# Towards In Vivo Wearable Diagnostics in Orthopaedics: Sensorized Bone Cement for Knee Spacer Applications

**DOI:** 10.64898/2026.04.21.719845

**Authors:** Nguyen Bach Tran, Laura Capogrosso, Christoph Dillitzer, Paul Morandell, Vincent Lallinger, Stephan Heller, Rainer Burgkart, Igor Lazic, Oliver Hayden

## Abstract

Periprosthetic joint infection (PJI) is a severe complication of total knee arthroplasty and is a leading cause of revision surgery, and is associated with significant morbidity. Two-stage exchange using antibiotic-loaded polymethylmethacrylate (PMMA) spacers remains the clinical gold standard, yet the decision to reimplant relies largely on indirect markers and clinical judgment, as no method allows continuous in situ assessment of infection resolution. Here, we report a sensorized knee spacer that transforms PMMA bone cement from a passive structural material into an active, wearable diagnostic device. A miniaturized multimodal sensor unit integrating optoelectronic and physicochemical sensing was embedded within the tibial spacer component and wirelessly coupled, enabling energy-efficient, 24/7 in vivo monitoring during the spacer interval for several months. We developed a reproducible encapsulation and integration strategy compatible with clinically realistic spatial, thermal, and mechanical constraints, without altering the established surgical workflow. The functionality of the embedded camera, spectrometer, and temperature sensors following cement integration was verified. Mechanical integrity and signal stability were confirmed under ISO-compliant dynamic biomechanical loading conditions. In vivo validation of the implantable wearable was preclinically demonstrated in a porcine model using human knee spacer dimensions. These findings establish the technical feasibility of sensor-integrated PMMA spacers and introduce bone cement as an enabling platform for smart orthopedic implants. Continuous, local monitoring of the peri-implant environment may open new pathways for evidence-based decision-making in infection management with temporary implantable wearables.

## Introduction

Due to demographic changes, the number of total joint arthroplasties (TJA) has risen significantly in recent years [1]. Osteoarthritis remains the leading cause for the implantation of hip and knee arthroplasties to restore the patient’s mobility [1, 2]. TJA procedures were performed on 4.3 million knees (primary TKA, 51.1%) and hips (primary THA, 32.4%) surgeries, with a patient average age of 65.6 years in 2024 in the US [3]. Despite excellent clinical outcomes of primary total knee arthroplasty (TKA), periprosthetic joint infection (PJI) represents one of the most severe complications, being the primary cause of early failure and revision surgeries in the United States [4]. The incidence of PJI following primary TJA implantation ranges from 0.3% to 32.5%, whereas the risk of infection increases significantly in revision [3, 5-10]. Furthermore, a 601% increase in demand for knee revision surgeries in the U.S. is projected between 2005 and 2030 [11]. Consequently, PJI poses a substantial economic burden, with estimated costs for revision TJAs in the U.S. market reaching approximately $2 billion by 2030 [12].

The two-stage revision procedure, including the use of a spacer, has become the gold standard for treating PJIs. In this approach, a spacer made of antibiotic-loaded bone cement serves as a temporary implant and placeholder until the permanent prosthesis can be implanted. Although this method continues to demonstrate high success rates, there is still no consensus on the optimal duration of the interim prosthesis, which typically ranges from two to six weeks [13, 14]. Currently, the interval between the two surgeries is determined solely on the basis of subjective clinical signs and low-specific serological markers of infection, such as swelling, redness, and elevated C-reactive protein (CRP) levels. [15]. Implementing a patient-specific decision-making tool during revision surgery has the potential to enhance patient outcomes. This approach advances antibiotic stewardship by limiting unnecessary antibiotic use and its associated adverse effects, while restricting length of stay to the medically necessary minimum, thereby reducing hospital resource utilization and associated costs.

## Background

Periprosthetic joint infection (PJI) exhibits substantial clinical heterogeneity, ranging from acute inflammatory presentations to low-grade, indolent courses characterized by subtle or nonspecific symptoms. This variability is reflected in biomarker profiles: commonly applied thresholds for CRP, erythrocyte sedimentation rate, synovial leukocyte count, and emerging molecular markers lack robustness across patients, pathogens, and stages of infection [16, 17]. Accordingly, fixed cut-off values, for example in clinical scoring systems, risk misclassification, as biomarker levels are modulated by patient age, immune response, comorbidities, surgical timing, and microbial virulence.

Longitudinal assessment of biomarker dynamics within the same patient therefore represents a more reliable diagnostic strategy. Tracking deviations from an individual baseline enables personalized interpretation of inflammatory changes and supports tailored diagnostic and therapeutic decision-making. However, enabling continuous (24/7) monitoring in vivo presents substantial engineering challenges. In particular, sensing functionality must be integrated into an antibiotic spacer while maintaining a design compatible with an accelerated regulatory approval pathway.

To realize an implantable, wearable diagnostic device, medical-grade biocompatible materials approved for in vivo use are required, alongside strict electrical isolation and the absence of direct contact between tissue or blood and electronic components. Additionally, a digitalized temporary knee implant (SmartSpacer) must be sufficiently rugged to withstand surgical handling, high mechanical loads during implantation, and intraoperative bleeding, all of which may impair sensor integrity or compromise optoelectronic measurements. Finally, continuous in vivo operation necessitates a power supply capable of supporting more than six weeks of monitoring at a clinically meaningful readout frequency while maintaining a practical shelf life.

The proposed implantable in vivo wearable device enables continuous access to biophysical parameters relevant to joint infection, facilitating objective and longitudinal assessment of the local infection status. The SmartSpacer allows real-time monitoring of the periarticular microenvironment without perturbing physiological conditions. Clinical evaluation is based on multimodal sensing, including local temperature, synovial membrane imaging, and spectroscopic measurements. A miniaturized camera and spectral sensor system enables precise spatiotemporal (4D) monitoring of potential PJI through non-invasive, synchronous acquisition of multiple biomarkers.

Integration of these multiparametric data streams enables patient-specific estimation of infection resolution, a critical consideration during two-stage exchange procedures. Real-time in vivo confirmation of successful antibiotic therapy may permit safer and earlier reimplantation of the definitive arthroplasty, with the potential to substantially reduce treatment duration and overall healthcare costs. Beyond its clinical impact, this approach provides deeper insight into the dynamic host-pathogen interaction and individualized treatment response.

A key enabler of the proposed sensing strategy is the observation that medical-grade polymethyl methacrylate (PMMA) bone cement without radiopaque additives is optically transparent. This property permits a range of optoelectronic sensing applications that have thus far not been exploited for diagnostic or theranostic purposes, while simultaneously minimizing regulatory risk by avoiding novel materials and preventing direct contact between sensors and biological tissue or blood. Prior work by Ray et al. (2019) and Veletic et al. (2022), there is considerable potential to integrate implants with sensors, thereby facilitating the measurement of diverse biomarkers and biosignals [18, 19]. Furthermore, Noordhuis et al. (2025) provided a comprehensive scoping review of the integration of biomedical sensors, with a particular focus on their application in hip and knee implants [20]. The patent under discussion delineates a prospective diagnostic and theragnostic application in total knee arthroplasty [21].

### Technical Aspect

The sensorized smart implant is equipped with a medically approved, integrated endoscopic camera on its anterior surface, enabling precise intra-articular synovial imaging. Additionally, the circuit board contains recesses that accommodate a miniaturized spectrometer unit (nanoLambda, South Korea), which is attached to the board and enables the measurement of spectral scattered-light reflection from the synovial membrane. To optimize energy efficiency, the spectrometer uses the camera’s integrated light source to capture the scattering spectrum. The SmartSpacer also includes three temperature sensors distributed across the circuit board to measure local circadian temperature at the site of infection. Digital temperature sensors are used to assess locally elevated temperatures associated with specific circadian rhythms, which are hallmarks of infection. Additionally, a position sensor has been integrated to detect the rotational movement of the SmartSpacer, consequently enabling the monitoring of the knee joint’s movement along the pitch and roll axes. The motion sensor plays a pivotal role in ensuring that images and spectra are captured at a fixed position and orientation of the circuit board. This is achieved by setting an angle-trigger event, which is a crucial component of the overall system. Data are transmitted wirelessly via Bluetooth Low Energy (BLE) to a mobile application accessible to the treating physician. All recorded sensor data are used with algorithm-based and statistical evaluation methods to differentiate between different bacterial concentrations. The electrical circuit board is composed of an FR4 composite material consisting of epoxy resin and glass fiber fabric, with a thickness of 1 mm. The circuit board dimensions are 55 × 34 mm (Figure 1A), and it has been specifically adapted to the dimensions of the smallest human spacer, 65 × 45 × 12 mm. In the SmartSpacer, the optoelectronic components and the circuit board are electrically isolated in epoxy resin and further encapsulated as an inlay within the PMMA bone cement of the spacer’s tibial component to prevent interactions with the surrounding tissue in the knee (Figure 1B). Only the endoscopic camera lens (Optasensor, Germany) is in contact with the synovial liquid with a 120° field of view. The PMMA dome functions as an optical interface, enabling light coupling for imaging with an endoscopic RGB CMOS 320×320-pixel sensor array camera and facilitating the detection of elastically scattered light along the extended optical path using a 390-1010 nm spectrometer with 5 nm resolution. Due to the volume of a human knee spacer, a standard PCB design was used, and flexible circuit board technologies for miniaturization were not required, thereby reducing additional risk for in vivo applications. The miniaturization and optimized arrangement of the optoelectronic components address the challenges posed by the epoxy coating’s encapsulation capability and the limited space available within the knee spacers, which are designed for both human medical use and animal studies (Figure 1C). The battery management system has been engineered to ensure a minimum shelf life of 2.5 years for the device. The incorporation of an additional magnet switch enables device deactivation via a magnet, thereby ensuring operational integrity. Moreover, this switch enables the device to be restored in the event of a malfunction, ensuring uninterrupted functionality.

**Figure 1:**
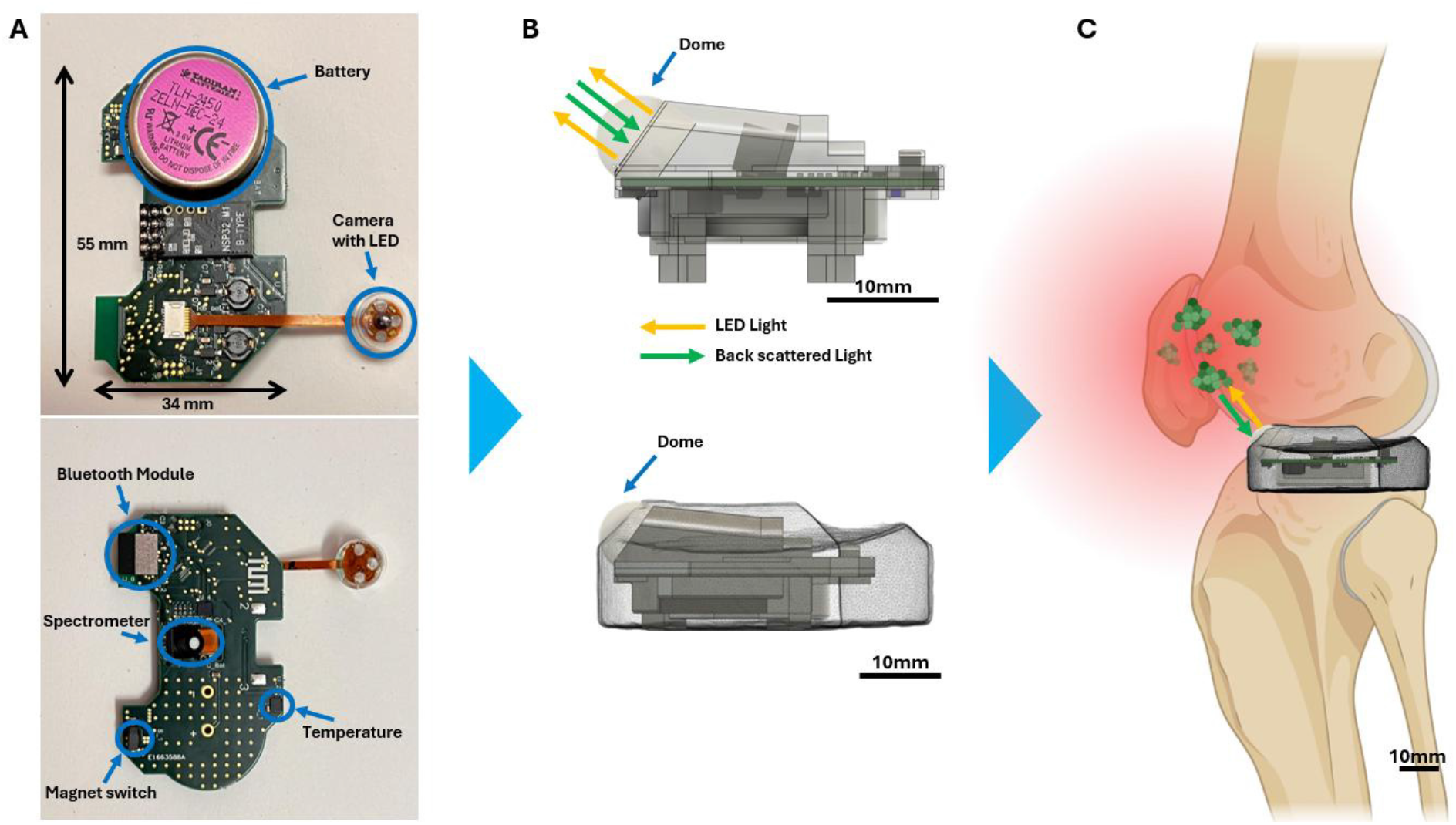
(A) The circuit board is equipped with a variety of sensors, and its dimensions are specified by the human tibia size. (B) The printed circuit board (PCB) will undergo a molding process comprising two encapsulation steps. The primary component is a medical-grade epoxy resin for electrical isolation and to prevent the formation of trapped air bubbles. The second encapsulation is that of PMMA bone cement with the medical-grade camera dome positioned towards the synovia. (C) The SmartSpacer is implanted after a functional check, without altering the surgical workflow.

### Preclinical Workflow

To verify and preclinically validate the diagnostic concept of in vivo wearables embedded in medical-grade PMMA, we used a commercially available human knee spacer from Heraeus bone cement [22], produced intraoperatively by the surgeon, and developed a tibial sensing unit that can be seamlessly incorporated into the clinical workflow. The surgical procedure in the preclinical pig trial is under general anesthesia with controlled ventilation and multimodal analgesia. Animals were positioned in left lateral decubitus, and the right knee was prepared and draped under sterile conditions. A lateral parapatellar approach was used with a 15 cm skin incision and arthrotomy, followed by subtotal resection of the infrapatellar fat pad and synovium, resection of the lateral meniscus, and proximal tibial resection (10 mm, 3° posterior slope). After anterolateral tibial subluxation, the medullary canal was prepared for the SmartSpacer keel. The SmartSpacer was implanted, and ligament balance was assessed dynamically; polymethylmethacrylate was added to address instability if observed. The joint was irrigated, hemostasis achieved, and the arthrotomy closed with resorbable sutures. Local infiltration anesthesia with ropivacaine was administered, followed by intra-articular inoculation with *Staphylococcus aureus* (initial dose 1 × 10^6^ CFU/mL in 1 mL saline). Wound closure was completed in layers, and animals were monitored for 5 ± 1 days with daily wireless SmartSpacer data acquisition and standardized clinical scoring.

During knee replacement surgery, the infected knee joint is excised, and the wound site is meticulously cleansed. The spacer is produced by the surgeon in the operating room, with the required length customized to the missing joint to ensure a precise fit. The surgeon commences the mixing of the bone cement at the operating table and places it in the holder. Following a 15-minute curing period [23], the spacer can be detached from the holder and implanted. In our use case, the pre-embedded board with the optics will be integrated into the bone cement spacer (Figure 1C).

In this study, an electronic device is embedded in bone cement using Heraeus Spacer mold sets for total knee arthroplasty. The implementation is verified by the functionality and biomechanical stability of the digitized sensor prototype. Following the embedding process, dynamic biomechanical testing was conducted in accordance with DIN ISO 14879-1 (2020), revealing that the SmartSpacer maintained its functionality after 300,000 cycles at 400 N. The initial in vivo test was conducted in a pig with a small human spacer (S) implanted in the right hind leg, which corresponds to the smallest human knee size, validating all SmartSpacer functionalities, including sensor readouts and wireless communication. The utility of biophysical biomarkers for longitudinally monitoring inflammation, infections, and bacterial eradication is part of the ongoing preclinical study.

## Materials and Methods

In the study, human S-spacers with a height of 12 mm, a medial-lateral width of 65 mm, and an anterior-posterior length of 45 mm were utilized as the target size. Using the smallest possible spacer dimension ensures the plate can be cast in any conceivable spacer size. The shaft facilitates robust and non-slip tibial fixation during implantation in the knee joint. The initial shaft length of 34 mm was reduced to 3 mm to align with the dimensions of the proximal tibia of the pig in the planned animal experiments and to minimize bone tissue loss. The underside of the spacer features a 20 × 20 mm base for the shaft. The bone cement utilized in this study, Palacos® R+G (Heraeus Inc., Germany), is characterized by its high viscosity and contains gentamicin as an antibiotic, as intended for human use. During the exothermic curing process, the temperature reaches 70-80°C without compromising the integrity of its embedded electronic components.

### Preparation to integrate the circuit board into bone cement

In order to ensure a successful integration of the PCB into the bone cement, it is imperative that three preparatory steps be followed meticulously.

It is imperative that the camera and the spectrometer be aligned to ensure optimal functionality, thereby maximizing the readout through the optical window. A 3D-printed bracket is employed to align the camera with the spectrometer. The camera is positioned at 30°, which is congruent with the intersection of the Heraeus mold. The angle to the spectrometer is dependent on the camera’s alignment with the synovial membrane, with a standard deviation of 19°. This configuration enables the spectrometer’s sensor chip, which is focused through one of the three apertures in the camera board, to detect the maximum possible backscattering of the camera light onto the membrane (Figure 2A). The utilization of 3D printing technology ensures the reproducibility of outcomes.

**Figure 2:**
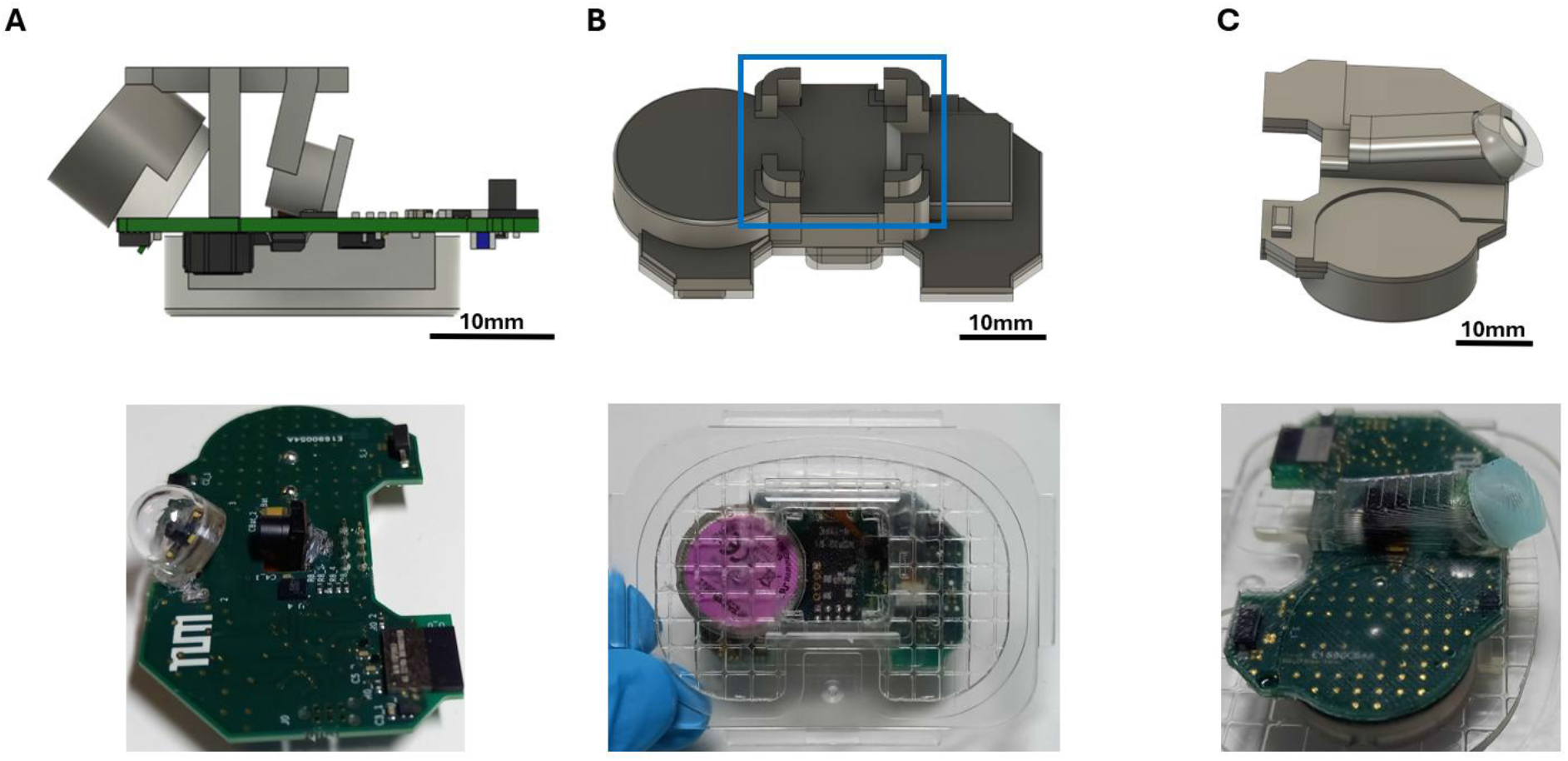
(A) demonstrates the use of a 3D-printed alignment tool to ensure the reproducibility of the angle between the camera and the spectrometer for all PCBs. (B) To ensure correct adhesion of the mold to the PMMA bone cement, it is essential to use a design click system that precisely centers and aligns the PCB, particularly the camera dome, with the synovial membrane. (C) It is imperative to note that during bone cement molding, the camera dome must be meticulously protected with a silicon cover. Following the removal of the surplus PMMA, the domes are rendered capable of providing unobstructed vision.

The subsequent step is to guarantee that the in-medical-graded covered PCB board is positioned within the spacer mold in such a manner that the bone cement is completely enclosed while the optical window remains accessible. The click system has been engineered to ensure that the circuit board is correctly positioned within the knee spacer and that the camera is against the Heraeus spacer’s wall, allowing a clear field of view following cementation. For this purpose, a four-legged click system is designed during the initial casting stage using epoxy resin. After the primary protective layer hardens, the circuit board can be attached to the Heraeus spacer cover (Figure 2B). Moreover, it is evident that this mounting bracket does not necessitate any alteration to the spacer set, thereby obviating the need for recertification of the system. Additionally, this will not alter the surgeon’s clinical workflow.

In order to prevent the occurrence of contamination of the optical window with bone cement during the casting process. A protective cap was designed using three-dimensional computer-aided design software, with the intention of its subsequent integration with the Heraeus spacer holder. This procedure is undertaken to prevent migration of bone cement over the camera dome, thereby ensuring the camera dome remains unobstructed (Figure 2C).

### Molding

During the manufacturing process of the SmartSpacer, the conventional method for filling the knee casting molds with bone cement was predominantly maintained. However, the spacer was initially embedded in epoxy resin. The preparation process was meticulously executed in two stages. Initially, a two-part pre-cast silicone mold was fabricated. The plate was then inserted into the silicone mold and subsequently encased in epoxy resin. The fabrication of snap-in holders on the precast component enabled attachment of the plate to the lid of the Copal® knee mold, which was subsequently encased in bone cement.

### Design

The design was conceived using the Fusion360 software (Autodesk, USA). The casting drawing was derived from the PCB and an uploaded CT scan of a fully cast spacer. The design was selected to ensure that the circuit board, with a layer thickness of approximately 0.5 mm, is isolated from the subsequent bone cement.

### Silicone impression

The configuration of the two-part silicone mold is instrumental in producing a comprehensive, bubble-free epoxy coating. The formation of bubbles during epoxy resin casting was significantly mitigated by continuous transitions, the reduction of narrow cross-sections within the mold, and the optimized placement of inflow and ventilation channels. Another salient point was to create a smooth surface finish on the epoxy layer to minimize friction and possible air pockets during casting with PMMA. Initially, the negative mold for the silicone mold was printed with a 3D printer using PLA filament (Bambu Lab, China) with a layer height of 0.1 mm. The component design corresponded to the circuit board design, with an additional epoxy coating. A rectangular frame, fabricated from PLA 3D printing with a thickness of 4.5 mm and a height of 4 mm, was affixed around the component to form a sealing lip, thereby enhancing the closure of the two silicone molds. The document was then affixed to the base of a separable container, with its halves connected by four M6 screws and nuts (Thorlabs, Germany). The container, measuring 87 × 148 × 46 mm, was fabricated from Siraya Tech’s gray Build Sonic Gray UV resin using the Anycubic Photon Mono X stereolithography 3D printer. The silicone was cast using a 1:1 ratio of components A and B of the green impression silicone (Laurenz+Morgan GmbH, Germany), a food-grade silicone. These components were weighed and mixed at room temperature. The initial phase of silicone mold fabrication involved meticulously weighing 50 g of each component. The two components were vigorously stirred in a beaker for 1 minute with a wooden spatula to ensure no air was introduced into the mixture. Subsequently, the silicone compound was deposited onto the PLA negative mold within the container and removed from the mold after 60 minutes of curing. The cured half of the silicone mold was rotated and positioned at the base of the container. The complete PLA negative mold, complete with inlet and ventilation channels (Figure 3A), was then inserted into the silicone mold. To establish a separating layer between the two silicone halves, the silicone surface was coated with approximately 5 ml of separating Vaseline (TFC Troll Factory) using a brush.

**Figure 3:**
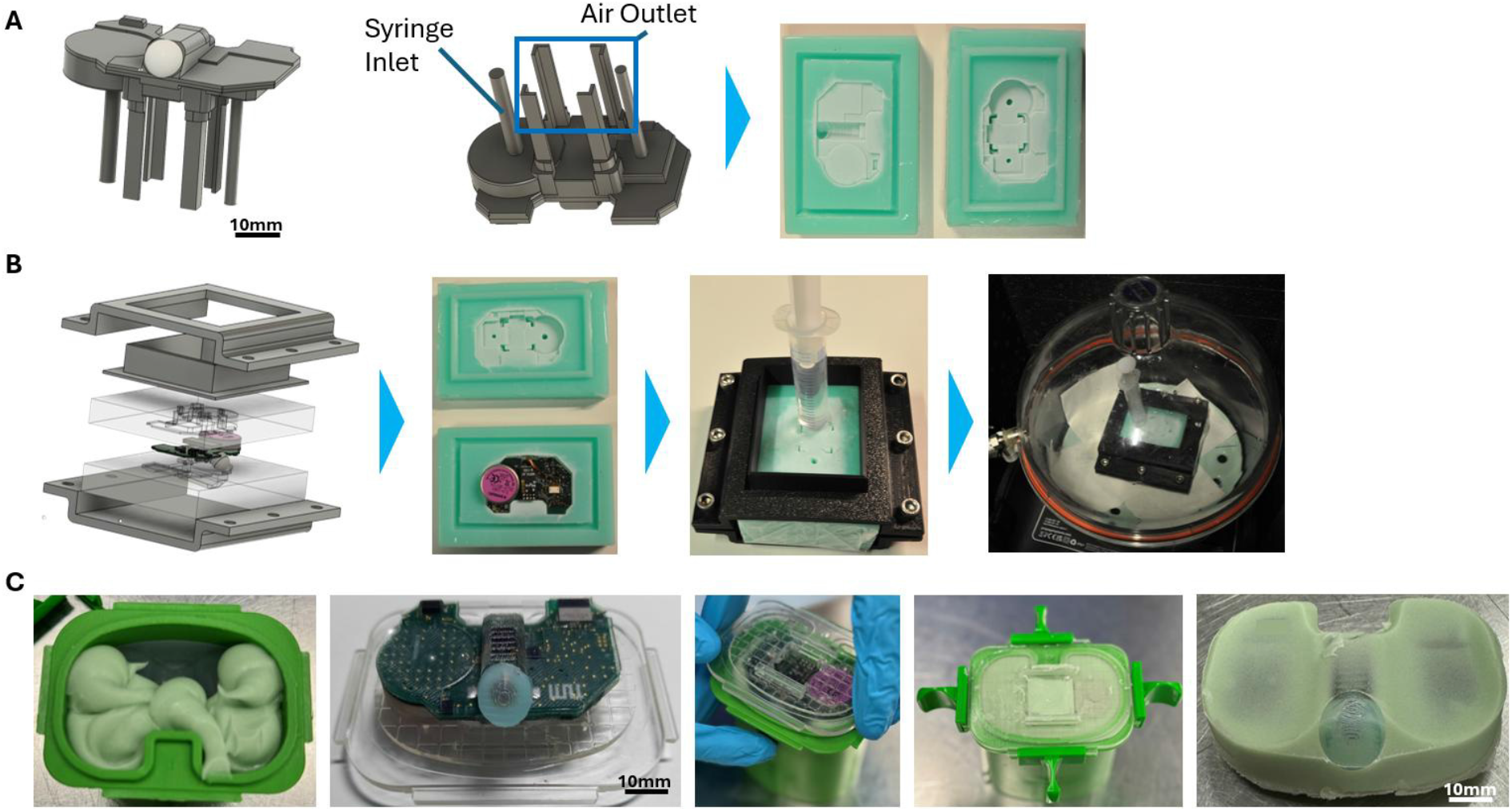
(A) The SmartSpacer is extended with an inlet for the purpose of introducing epoxy resin, in addition to multiple outlets for the purpose of creating a silicone mold. (B) The printed circuit board (PCB) is to be positioned between the silicon mold and secured using 3D-printed shells. (C) illustrates the insertion of the epoxy-resin-encapsulated PCB, the camera dome cover, and the click system. The primary workflow of the procedure will remain unaffected.

For the second silicone half, 75 g of each of the two silicone components was weighed and, after mixing, poured onto the first silicone half.

Concurrently, the negative mold of the pre-cast was secured in the intended position by a metal weight. After 60 minutes, the second silicone half could be extracted from the container and separated with ease from the first silicone half, owing to the separating vaseline (Figure 3A).

### Epoxy encapsulation of the circuit board

The epoxy resin EPO-TEK® MED-301 (Epoxy Technology Inc., Massachusetts, USA), which has received medical approval, was utilized for the epoxy coating. The product exhibits low viscosity and a pot life of 45 minutes. However, the required curing time varies with ambient temperature. In this process, a curing time of 16 hours at 37°C was selected. In preparation for casting, the circuit board was initially positioned in the silicone mold and then enclosed in the secondary silicone half. The sealing lip of the silicone casting mold enabled the connection of both halves in a form-fitting manner. The two silicone molds were meticulously sealed along the sides with transparent adhesive tape, ensuring that the inlet and ventilation channels on the top remained unobstructed. A 2 mm-thick rectangular template, fabricated from PLA, was affixed to the upper surface of the connected silicone mold. This was done to restrict the epoxy resin from escaping from the outlet channels. The template was adhered to a 5 mm-thick two-part PLA fixture. The two-piece fastener ensured that the two silicone halves were evenly fixed in place, thus preventing the epoxy resin from escaping. The two fastener components were affixed along the longer edges using a total of six M6 screws and nuts. Both the template and the two-piece holder were printed with a nozzle diameter of 0.8 mm and a layer thickness of 0.2 mm (Figure 3B). The epoxy resin and hardener were initially combined at a ratio of 1:4. In the production of the epoxy pre-coating for a circuit board, a quantity of 11.25 g of epoxy resin and 33.75 g of hardener were utilized. The two components were stirred for 2 minutes, forming a homogeneous mixture. Subsequently, the epoxy resin mixture was vacuum-degassed in a desiccator at 0.9 bar for 5 minutes. Two 12 ml disposable syringes, equipped with Luer slip attachments, were each filled with 10 ml of the epoxy mixture. The syringes were then filled to 12 ml, creating an internal air space. To eliminate any residual air bubbles, the two syringes were positioned vertically in the desiccator, with the cones oriented upwards, using a cardboard holder. The syringes were then degassed under vacuum for 5 minutes. Subsequently, the first syringe was vented and completely injected into the closed silicone mold via the inlet’s opening. The epoxy resin was observed to be extruding from all five outlet channels. After venting the second syringe, approximately 3 ml of its contents were injected into the silicone mold, with the syringe left in the mold. The sealed silicone mold, with the syringe positioned on top, was placed in the desiccator for the third vacuum cycle. The mold was then vacuumed for an additional five minutes. After removing the silicone mold from the desiccator, an additional 5 ml of the mixed solution was injected into the silicone. The final assembly was then placed in a drying oven at 37°C for 16 hours to achieve complete curing. The exothermic reaction that transpired during this process resulted in the observation of heat development within the range of 30 to 60°C. The epoxy-coated circuit board was then extracted from the mold by separating the PLA template, which was bonded to an epoxy layer, from the silicone mold using a small hand hacksaw. This procedure was preceded by loosening the screws of the two-piece fastener. The template was designed to prevent direct contact between the liquid epoxy and the two-piece holder. Subsequently, the epoxy-coated circuit board was extracted from the silicone mold and reworked using side cutters, for example, to remove any residual material from the inlet and outlet channels.

### SmartSpacer molding

The fabrication of native, fully cast spacers in size S involved the use of Copal® knee molds, which were filled with Palacos® R+G bone cement at approximately 20°C. In the context of the Copal® knee molding procedure, the utilization of the sterile disposable mold was confined exclusively to the tibial component, inclusive of the associated cover and shaft.

The tibial component is height-adjustable, allowing the tibial plateau height to be flexibly adjusted over a range of 12 mm to 37 mm. The molds are composed of medical-grade plastic and were used multiple times during the manufacturing process, provided that their structural integrity and functional capabilities remained unaltered. In preparation of the molds, the shaft on the lid was reduced to a length of 12 mm to ensure the specified length of the tibial shaft. To produce a PMMA spacer with a minimum number of air pockets, the bone cement is mixed under vacuum. For this purpose, the Palamix® UNO cartridge vacuum mixing and application system (Heraeus Inc., Germany) was used according to the manufacturer’s instructions. The mixing system comprised a cartridge, a mixing rod with an integrated mixing paddle, an application snorkel, a vacuum hose, and a funnel. The Palamix® vacuum pump, powered by compressed air, was used in conjunction with a hospital-compressed air connection. This resulted in a 5 bar vacuum within the mixing system cartridge. The Palamix® cement gun is also a necessary accessory for producing tibial plateaus.

The SmartSpacers were produced using a standardized procedure. This procedure was implemented to minimize segregation and air pockets while ensuring the best possible workability of the bone cement. Prior to use, both the polymer powder and the monomer liquid were cooled in a refrigeration unit set to 4°C for at least 24 hours. This slight reduction in viscosity of the bone cement was intended to optimize processing and extend the processing and curing times [23, 24]. The objective was to achieve a homogeneous polymer network with a reduced number of voids for the tibial plateau. The two components were removed from the refrigerator immediately before being poured into the Palamix® uno vacuum mixing system, with a maximum lead time of 15 min. Initially, the height of the tibial component of the Copal® knee mold was set to 12 mm. Subsequently, the lid, which featured a reduced shaft, was positioned over the mold and secured using the four slide locks. In this section, the SmartSpacer is described in relation to the PCB, which is positioned centrally within the cement injection hole, with the mounting system from the epoxy resin coat in place.

The two cement components were subsequently introduced into the mixing system by filling the cartridge through the funnel, which was positioned on a fixture within the packaging. Initially, 40 ml of the monomer liquid was decanted, followed by the incorporation of 81.6 g of the polymer powder, yielding two spacers using a mixing cartridge. This prevented the formation of powder pockets, as polymerization began immediately at the surface. After sealing the mixing cartridge, a vacuum was applied for 10 s.

Subsequent to the complete hardening of the bone cement, which occurred approximately 9 to 10 min following the mixing of the two components, the tibial plates were retrieved from the molds. The final sTJA in the fabrication process involved removing burrs and unevenness using grinding and deburring tools, ensuring a smooth, polished finish. In general, when manufacturing PMMA spacers, a storage time of 24 to 48 hours is required to ensure complete polymerization (Figure 3C).

### Verification

#### Dynamic biomechanical testing

The present study sought to ascertain the mechanical fatigue strength of the SmartSpacer. To this end, dynamic fatigue tests were performed in accordance with ISO 14879-1 (2020). The servo-hydraulic testing machine was utilized to conduct the aforementioned tests. The experimental apparatus utilized for this investigation comprised the Amsler HC10 load frame (ZwickRoell, Ulm, Germany) and the testXpert R testing software, version V2.0.0 (ZwickRoell GmbH Co. KG, Ulm, Germany). The configuration enabled the loading device to be positioned at an elevated level, thereby facilitating control of the test cylinder within the testing machine (Figure 4). In the course of the dynamic tests, the spacers were subjected to cyclic loading with a maximum force of 400 N and a minimum force of 40 N. To this end, a sinusoidal dynamic load waveform at 2.5 Hz was applied. According to ISO 14879-1 (2020), the minimum force should be 10% of the maximum force, and the load frequency should not exceed 10 Hz [25]. In the context of the dynamic test, the parameters originally designed for a permanent implant underwent adaptation for a temporary implant in this experiment. Instead of the 10 million cycles stipulated in the test standard for permanent implants, the test duration was reduced to 300,000 cycles. This number of cycles is designed to simulate human gait for approximately three months. This adjustment was made in consideration of the tibial knee spacer’s maximum retention period of six weeks during clinical utilization. Moreover, a relatively high test frequency of 2.5 Hz was selected to reduce the testing duration. However, it should be noted that the physiological movements of a patient with a temporary implant are approximately 1 Hz. The maximum load of 900 N specified in ISO 14879-1 (2020) was significantly reduced for this test series, as the knee spacer molded from the Copal® knee molds is designed exclusively for patients who use mobility aids such as crutches throughout the entire implantation period and therefore do not place full weight on the knee joint. Furthermore, the unilateral free-swinging load on the tibial component delineated in the standard constitutes an extreme load case that does not accurately reflect physiological movement. Consequently, this value was employed as the maximum load for the test series. The SmartSpacer was exposed to the load following a storage period of 20 to 29 days in a light- and moisture-protected environment at 20 to 24°C, with the force applied to the left side of the tibial component. The tibial plateaus were packaged in sealed plastic bags. The tests were terminated promptly upon the attainment of the stipulated deformation or force limits of 500 N.

**Figure 4:**
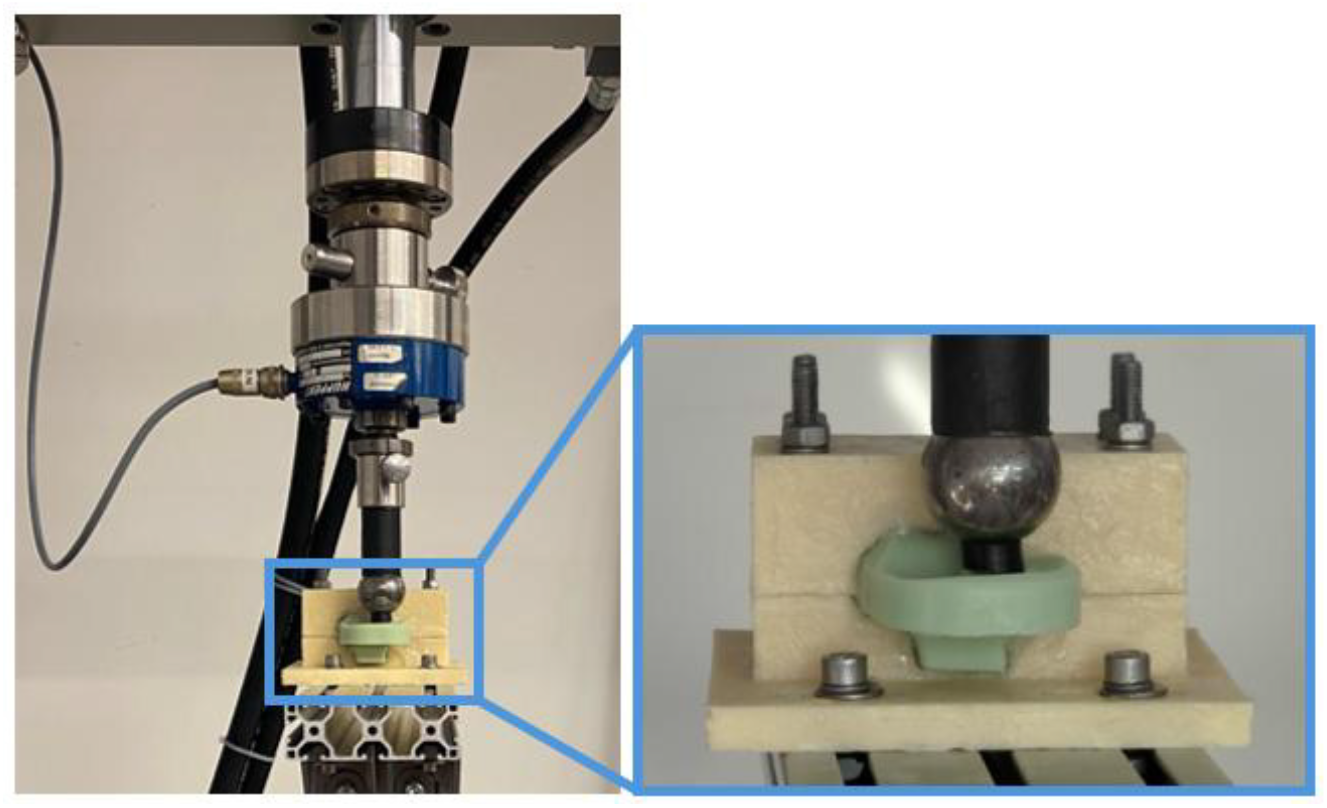
Setup of the biomechanical testing device. The biomechanical load structure is delineated in accordance with ISO standards.

**Figure 5:**
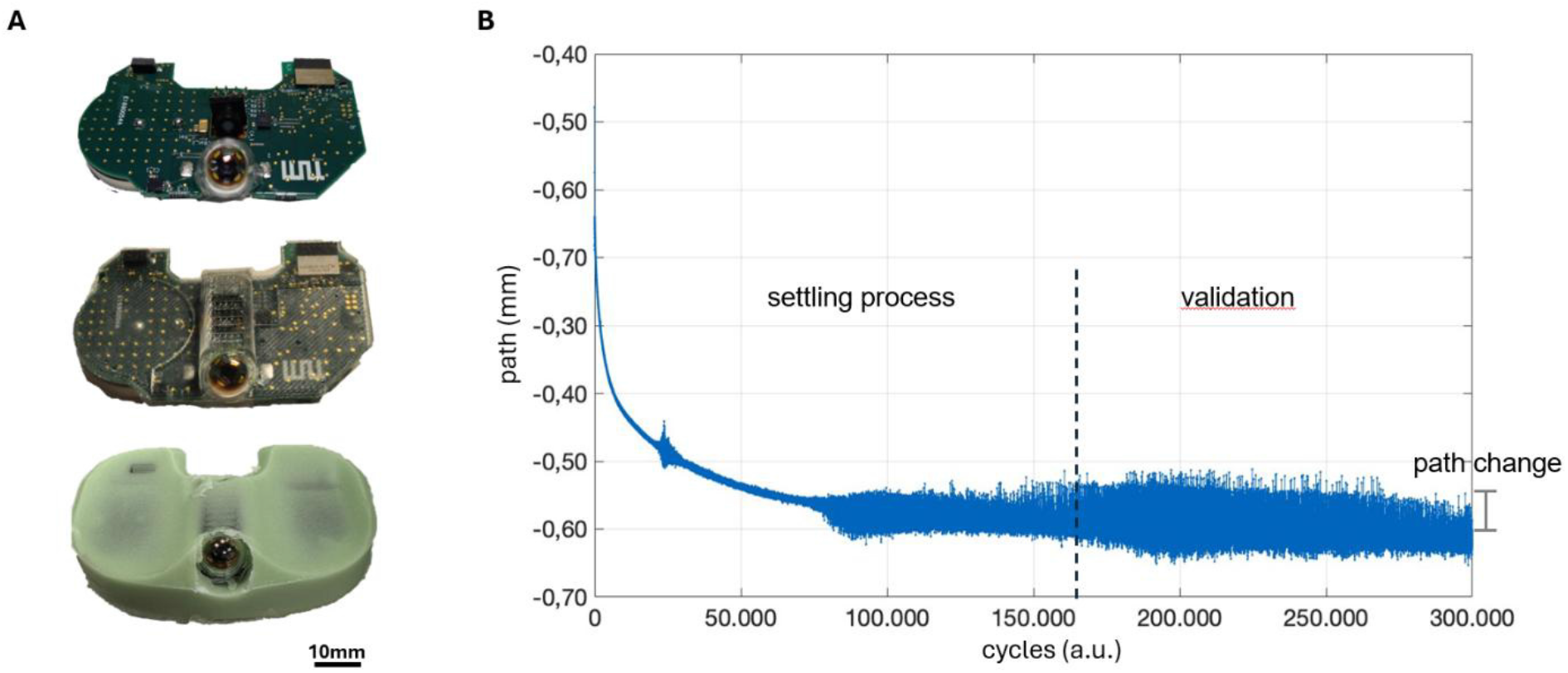
(A) Shows the circuit board on top without, then with epoxy resin, and final with the PMMA encapsulation. (B) The biomechanical dynamic testing shows the progression of change-of-direction after over 300.000 cycles.

### In vivo testing

The SmartSpacer was implanted in a pig’s knee (n=1) and tested for functionality for 24 hours (Figure 4B). The rationale behind the selection of the pig as a subject for this experiment is twofold. Firstly, its knee joint anatomy is highly analogous to that of humans. Secondly, its immune response system for later infection monitoring is similar to that of humans. However, the latter will not be discussed here. The pig weighed more than 100 kg. This weight is necessary because the pig’s knee corresponds to the size of the human S spacer. The surgical TJA followed the procedure used for total knee replacement in humans. The data were recorded at various intervals: images every hour, spectra every 10 minutes, and temperatures every 10 minutes.

## Results

The procedure delineated in the Methods section demonstrates that the circuit board is entirely encapsulated with EPOTEK epoxy resin (Epoxy Technology Inc., Massachusetts, USA). In addition to insulating the electronics from the bone cement, the encapsulation technique also creates the special click-in system. The removal of the optical window from the overflowing epoxy resin was performed cleanly and without damage, facilitated by the release coating. This facilitated an unobstructed view of the imaging and spectrum recording. Furthermore, the injection molding process with bone cement and the embedded circuit board module also achieved successful results. The circuit board, which had been treated with an additional epoxy coating, was completely encased by the bone cement without any discernible air pockets. The compact dimensions of the circuit board module, in conjunction with the miniaturization of the optoelectronic components, facilitated the encapsulation of the spacer with PMMA, even with the circuit board in place. The use of a silicone encapsulation cap ensured a clear window for the camera on the spacer’s front (Figure 4A).

### Dynamic biomechanical testing

For dynamic testing, the SmartSpacer was loaded on the sensor side (left half) to a maximum force of 400 N for 300,000 cycles. After loading, no cracks were detected. The circuit board spacer demonstrated uninterrupted functionality throughout testing, successfully transmitting all sensor data. Notwithstanding the 33-hour test duration, the device maintained a persistent connection to the software. The distance traversed by the pressure stamp during the cyclic loading process is represented by a sinusoidal vibration curve, with the upper and lower reversal points delineated by two envelopes. The envelope curve for the lower reversal points of the displacement as a function of the cycles is illustrated, utilizing the dummy spacer as a representative example. As illustrated in Figure 4B, the envelope curve of the lower reversal points of the displacement is depicted as a function of the cycles during the dynamic test of the dummy spacer. The curve illustrates the deformation of the specimen during cyclic loading and shows negative values, as the compressive load on the spacers is applied in the negative z-direction. The curve indicates that the displacement only stabilizes after approximately 130,000 to 170,000 cycles. During the initial cycles, a so-called “settling-in process” can be observed, as the sample must first stabilize in the holder due to the additional fixation. Consequently, an interval of 170,000 to 300,000 cycles was considered for the analysis of the elastic behavior. In this case as well, the change in sample displacement over a specific time interval was analyzed to evaluate the results. A meticulous examination of the test specimens was conducted to detect any potential cracks.

### In vivo testing

Subsequent to the implantation of the SmartSpacer in the pig’s knee (Figure 6A), data were collected over a period of 8 hours. The camera’s view captures the synovial membrane located posterior to the patella. However, due to postoperative bleeding, the initial visual impression is a black-to-very dark image, making it challenging to discern with the naked eye. As time progresses, the bleeding diminishes, and the structure of the synovial membrane becomes increasingly discernible (Figure 6B). The recorded backscatter spectrum displays the fundamental spectrum of the white-light LED, emitted from the camera board to the membrane and subsequently backscattered through the holes on the camera board to the spectrometer unit (Figure 6C). With time, the spectral intensity becomes discernible, enabling the estimation of bacterial concentrations. The temperature curve demonstrated an average temperature of 37.8°C, which corresponds to the standard body temperature post-surgery. (Figure 6D). The average temperature is measured using three temperature sensors distributed across the PCB to cover the entire area of the spacer.

**Figure 6:**
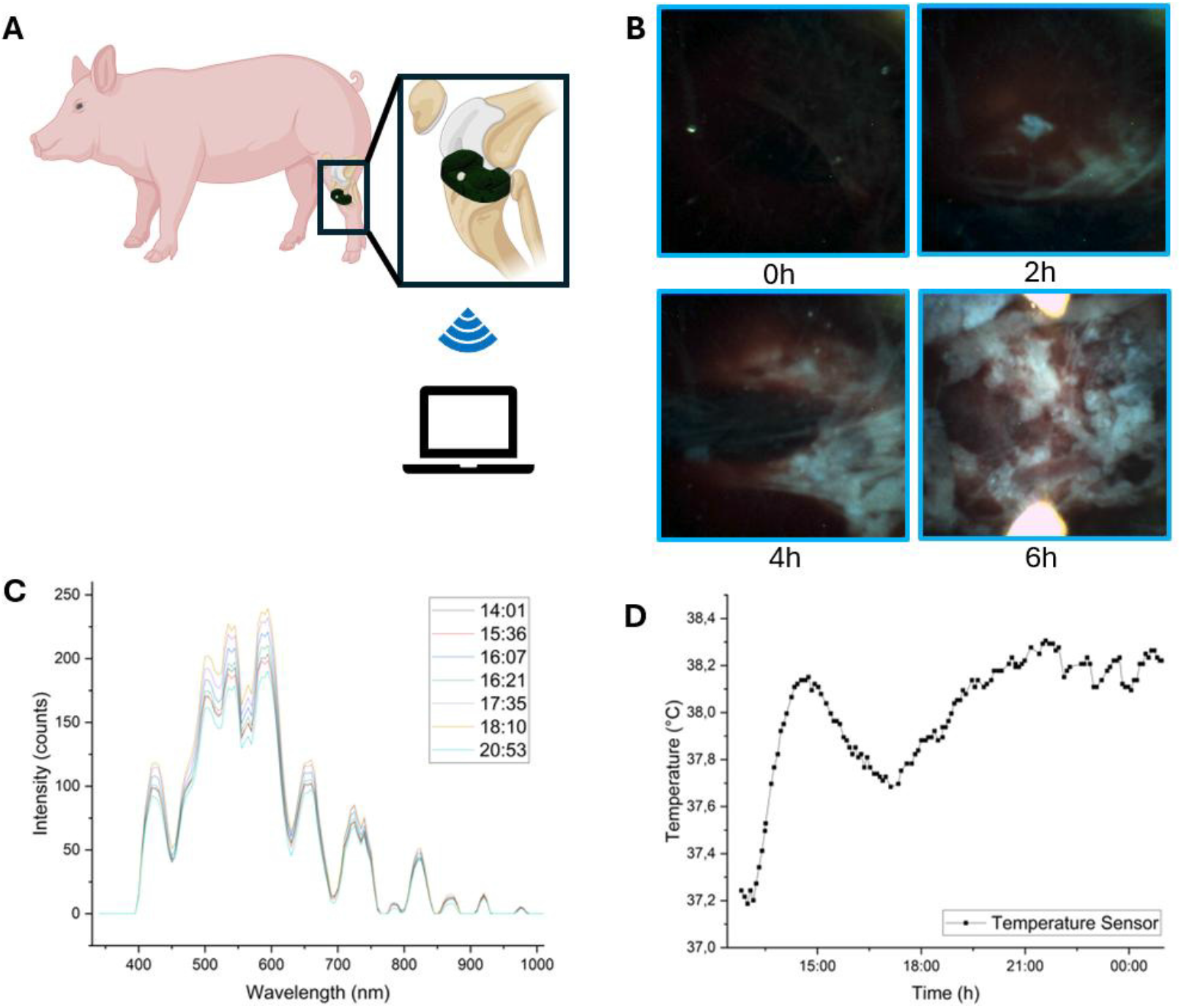
(A), In vivo pig trial with the sensorized spacer. (B), The camera is configured to capture images at hourly intervals after the surgical intervention. After 6 hours, the images became significantly clearer, attributable to the cessation of bleeding. (B) It has been demonstrated that spectra exhibiting divergent intensity counts can be discerned. (C) It is possible to monitor the temperature sensor. The plot illustrates the mean value of three PCB-installed temperature sensors. The patient’s temperature is rising, reflecting a trauma phase after the implantation with fever.

## Discussion

The work presented here demonstrates the feasibility of integrating optoelectronic sensors into a knee spacer for an implantable wearable. The primary objective of the initial TJA, which involved immersing the circuit board in epoxy resin, was to create a barrier between the electronics and the bone cement. This TJA also functioned as a protective covering, thereby preventing contact with tissue in the event of a fracture in the bone cement or leakage from the battery. During vacuuming, it was not always possible to completely remove all air bubbles from the casting. This phenomenon resulted in the formation of cavities at random locations within the casting. To optimize this process, standard operating procedures must be implemented. The second embedding of the circuit board into the bone cement demonstrated the feasibility of fully integrating it into the bone cement during the initial casting. The click-in system demonstrated its efficacy by firmly affixing the circuit board to the Heraeus mold at a predetermined distance. This ensured that the optical window was not compressed against the spacer mold wall, thereby preserving its structural integrity. The silicone cap on the dome ensured that the lens remained scratch-free and free of bone cement residue. A thin PMMA layer was observed to encapsulate only the magnetic switch located on the posterior side of the spacer. Degradation of this PMMA layer over a six-week implantation period may lead to partial exposure of the magnetic switch. However, the component is additionally protected by a medically approved epoxy encapsulation, mitigating direct tissue contact. Nevertheless, optimization of the magnetic switch placement may be considered in future design iterations to further enhance robustness. Future work will also focus on validating the proposed biomarkers through multimodal sensor fusion.

## Statements and Declarations

### Data availability statement

The data that support the findings of this study are available from the corresponding author upon request. Additional detailed results will be reported in a forthcoming preclinical study.

### AI tool usage

AI tools, such as DeepL and Microsoft Copilot, were used exclusively to enhance linguistic precision and clarity.

## Author Contribution

R. Burgkart, L. Capogrosso, and N. B. Tran conceived the experiment(s). L. Capogrosso, V. Lallinger, I. Lazic, P. Morandell, and N. B. Tran conducted the experiments. N. B. Tran analyzed the results and drafted the manuscript. S. Heller, C. Dillitzer, I. Lazic, R. Burgkart, and O. Hayden edited the manuscript.

## Acknowledgement

The author thanks Dr. J. Werner, Dr. J. Reiser from TUM University Hospital - Center for Preclinical Research, and Dr. A. Obermeier. Dr. Werner and Dr. Reiser helped with the in vivo study. Dr. Obermeier support the biomechanical testing. The authors thank the BMFTR (Funding number: 01EK2107A, 01EK2107B) for the financial support.

## References

[1] D. C. Wirtz, H. Reichel, G. Matziolis, and T. Pfitzner, Endoprothetik des Kniegelenkes, 2 ed. Springer: Springer Berlin, Heidelberg, 2023, p. 423.

[2] C. Otto-Lambertz, A. Yagdiran, F. Wallscheid, P. Eysel, and N. Jung, “Periprosthetic Infection in Joint Replacement,” Dtsch Arztebl Int, vol. 114, no. 20, pp. 347–353, May 26 2017, doi: 10.3238/arztebl.2017.0347.

[3] C. N. Carender, V. Hegde, B. R. Levine, J. I. Huddleston, 3rd, and A. Cohen-Rosenblum, “Highlights of the 2024 American Joint Replacement Registry Annual Report,” Arthroplast Today, vol. 33, p. 101727, Jun 2025, doi: 10.1016/j.artd.2025.101727.

[4] K. J. Bozic et al., “The epidemiology of revision total knee arthroplasty in the United States,” Clin Orthop Relat Res, vol. 468, no. 1, pp. 45–51, Jan 2010, doi: 10.1007/s11999-009-0945-0.

[5] M. M. Alrayes and M. Sukeik, “Two-stage revision in periprosthetic knee joint infections,” World J Orthop, vol. 14, no. 3, pp. 113–122, Mar 18 2023, doi: 10.5312/wjo.v14.i3.113.

[6] S. M. Kurtz, K. L. Ong, E. Lau, K. J. Bozic, D. Berry, and J. Parvizi, “Prosthetic joint infection risk after TKA in the Medicare population,” Clin Orthop Relat Res, vol. 468, no. 1, pp. 52–6, Jan 2010, doi: 10.1007/s11999-009-1013-5.

[7] S. M. Mortazavi, J. Molligan, M. S. Austin, J. J. Purtill, W. J. Hozack, and J. Parvizi, “Failure following revision total knee arthroplasty: infection is the major cause,” Int Orthop, vol. 35, no. 8, pp. 1157–64, Aug 2011, doi: 10.1007/s00264-010-1134-1.

[8] J. R. Palmer, T. S. Pannu, J. M. Villa, J. Manrique, A. M. Riesgo, and C. A. Higuera, “The treatment of periprosthetic joint infection: safety and efficacy of two stage versus one stage exchange arthroplasty,” Expert Rev Med Devices, vol. 17, no. 3, pp. 245–252, Mar 2020, doi: 10.1080/17434440.2020.1733971.

[9] G. Peersman, R. Laskin, J. Davis, and M. Peterson, “Infection in Total Knee Replacement - A Retrospective Review of 6489 Total Knee Replacements,” Clinical Orthopaedics & Related Research, vol. 392, pp. 15–23, 2001. [Online]. Available: https://www.ovid.com/jnls/clinorthop/fulltext/00003086-200111000-00003∼infection-in-total-knee-replacement-a-retrospective-review.

[10] J. E. Phillips, T. P. Crane, M. Noy, T. S. Elliott, and R. J. Grimer, “The incidence of deep prosthetic infections in a specialist orthopaedic hospital: a 15-year prospective survey,” J Bone Joint Surg Br, vol. 88, no. 7, pp. 943–8, Jul 2006, doi: 10.1302/0301-620X.88B7.17150.

[11] S. Kurtz, K. Ong, E. Lau, F. Mowat, and M. Halpern, “Projections of primary and revision hip and knee arthroplasty in the United States from 2005 to 2030,” J Bone Joint Surg Am, vol. 89, no. 4, pp. 780–5, Apr 2007, doi: 10.2106/JBJS.F.00222.

[12] C. Lavernia, D. J. Lee, and V. H. Hernandez, “Theincreasingfinancialburdenofkneerevisionsurgery in the United States,” Clin Orthop Relat Res, vol. 446, pp. 221–6, May 2006, doi: 10.1097/01.blo.0000214424.67453.9a.

[13] H. M. L. Muhlhofer et al., “Synovial aspiration and serological testing in two-stage revision arthroplasty forprostheticjoint infection: evaluationbefore reconstruction with amean follow-up of twenty seven months,” Int Orthop, vol. 42, no. 2, pp. 265–271, Feb 2018, doi: 10.1007/s00264-017-3700-2.

[14] E. M. Schwarz et al., “2018 International Consensus Meeting on Musculoskeletal Infection: Research Priorities from the General Assembly Questions,” J Orthop Res, vol. 37, no. 5, pp. 997–1006, May 2019, doi: 10.1002/jor.24293.

[15] R. von Eisenhart-Rothe and R. Burgkart, “Estimates of case numbers for TEP revisions in Germany for health economic evaluation of smart joint spacer.,” ed. Klinik fOr Orthopadie TU MOnchen, 2019.

[16] G. Solarino, D. Bizzoca, L. Moretti, G. Vicenti, A. Piazzolla, and B. Moretti, “What’s New in the Diagnosis of Periprosthetic Joint Infections: Focus on Synovial Fluid Biomarkers,” Trop Med Infect Dis, vol. 7, no. 11, Nov 7 2022, doi: 10.3390/tropicalmed7110355.

[17] K. Tsikopoulos and G. Meroni, “Periprosthetic Joint Infection Diagnosis: A Narrative Review,” Antibiotics (Basel), vol. 12, no. 10, Sep 27 2023, doi: 10.3390/antibiotics12101485.

[18] T. R. Ray et al., “Bio-Integrated Wearable Systems: A Comprehensive Review,” Chem Rev, vol. 119, no. 8, pp. 5461–5533, Apr 24 2019, doi: 10.1021/acs.chemrev.8b00573.

[19] M. Veleticet al., “Implants with Sensing Capabilities,” Chem Rev, vol. 122, no. 21, pp. 16329–16363, Nov 9 2022, doi: 10.1021/acs.chemrev.2c00005.

[20] P. H. H. Noordhuis, P. C. Jutte, A. G. P. Kottapalli, C. J. C. Lamoth, and C. C. C. Roossien, “Advancements in Biomedical Sensors for Early Detection of Failure in Hip and Knee Implants: Scoping Review on Potential Sensors for Implant Integration,” Ann Biomed Eng, vol. 53, no. 10, pp. 2392–2407, Oct 2025, doi: 10.1007/s10439-025-03780-5.

[21] R. Burgkart, O. Hayden, A. Obermeier, and R. von Eisenhart-Rothe, “EUROPEAN PATENT APPLICATION - THERAGNOSTIC ENDOPROSTHETIC SPACER,” Germany Patent EP 3 815 649 A1, 2019.

[22] H. Medical. “Copal® knee moulds. Heraeus Medical. [Gebrauchsanweisung] 2019.” https://www.heraeus-medical.com/de/healthcareprofessionals/products/copal-knee-moulds (accessed 11, 2024).

[23] K.-D. KOhn, PMMACements. Springer, 2014, p. 291.

[24] M. Arora, E. K. Chan, S. Gupta, and A. D. Diwan, “Polymethylmethacrylate bone cements and additives: A review of the literature,” World J Orthop, vol. 4, no. 2, pp. 67–74, Apr 18 2013, doi: 10.5312/wjo.v4.i2.67.

[25] D. Media. “ISO 14879-1:2020-07 Implants for surgery - Total knee-joint prostheses - Part 1: Determination of endurance properties of knee tibial trays.” https://www.dinmedia.de/de/norm/iso-14879-1/327849071 (accessed 03, 2026).

